# Spatially localized expression of glutamate decarboxylase *gadB* in *Escherichia coli* O157:H7 microcolonies in hydrogel matrix

**DOI:** 10.1101/2023.02.16.528592

**Authors:** Cédric Saint Martin, Nelly Caccia, Maud Darsonval, Marina Grégoire, Arthur Combeau, Grégory Jubelin, Florence Dubois-Brissonnet, Sabine Leroy, Romain Briandet, Mickaël Desvaux

## Abstract

Functional diversity within isogenic spatially organized bacterial populations has been shown to trigger emergent community properties such as stress tolerance. Taking advantage of confocal laser scanning microscopy combined with a transcriptional fluorescent fusion reporting at single cell scale the expression of the glutamic acid decarboxylase *gadB* in *E. coli* O157:H7, it was possible to visualize for the first-time spatial patterns of bacterial gene expression in microcolonies grown in a gelled matrix. The *gadB* gene is involved in *E. coli* tolerance to acidic conditions and its strong over-expression was observed locally on the periphery of embedded microcolonies grown in acidic hydrogels. This spatialization of *gadB* expression did not correlate with live/dead populations that appeared randomly distributed in the colonies. While the planktonic population of the pathogens was eradicated by an exposition to a pH of 2 (HCl) for 4h, mimicking a stomachal acidic stress, bacteria grown in gel-microcolonies were poorly affected by this treatment, in particular in conditions where *gadB* was spatially overexpressed. Consequences of these results for food safety are further discussed.

## 1) Introduction

*Escherichia coli* is a commensal bacterium found in the gut of mammals that plays an integral part in the digestive process. However, some strains of *E. coli* are pathogenic and represent a public health issue when they reach the production lines of food industries. Besides obvious evisceration accidents contaminating the meat at the slaughter state, food contamination of animal products (meat and milk products), vegetable or water usually occur through direct or indirect fecal contamination ^1–3^. Storage conditions and holding temperature are then major contributors to bacterial growth and survival in food products ^4^. Shigatoxin (Stx) encoding *E. coli* (STEC) are the third most common foodborne zoonosis in Europe ^5^ and amongst STEC, the serotype O157:H7 is a commonly identified agent in patients. *E. coli* O157:H7 are enterohaemorrhagic *E. coli* (EHEC) responsible for bloody diarrhea when the intestinal lining is broken by the presence of Stx. A possible outcome of Stx passing in the bloodstream is damage to the kidneys that can lead to a hemolytic uremic syndrome (HUS), which itself lead to fatal outcomes in 5% of cases ^6^. Children are especially at risk and *E. coli* O157:H7 is still the main cause of pediatric HUS ^7^. At the level of the European union, regulations ask for the absence of this pathogen in 25g of germinated seeds (Regulation CE 209/2013, amendment 2073/2055), but no equivalent exist for meat products. Precautionary measures for meats exist at a state level in the Union (France, DGAL/SSDSA/2016-353). However, the pathogen is still routinely detected at levels above 100 CFU/g in more than 1% of all red meats, the main vector of infection for this pathogen ^5,8^.

Structured food matrices are a continuity of heterogeneous local microenvironments harboring multiple micro-gradients that can evolve with time and microbial activity ^9^. This leads bacterial cells in food matrices to face different biotopes in which their growth and behavior can diverge from observations in liquid laboratory media. Therefore, environmental conditions of food matrices can prompt high phenotypic diversity in microbial populations as the cells adapt to local microenvironments ^10,11^. In comparison with their planktonic counterparts, phenotypic diversity in structured communities can influence bacterial fitness and behavior, such as increase expression of virulence genes ^12^, higher tolerance to antimicrobials agents and thermal stress ^13,14^, or improved cell motility ^15^. While several studies reported emergent properties of bacterial community in food matrices at the population level ^13,16–19^, no experimental evidence has yet been reported on the spatial heterogeneity of gene expression at the scale of single cells.

The stomachal phase after food ingestion exposes bacteria to strong acidic pH conditions for several hours and is credited for the highest population reduction of the bacterial load. High tolerance to acidic conditions is therefore necessary for foodborne pathogens, and involved systems that regulate intracellular pH. The glutamic acid decarboxylase (GAD) is one among various systems of acid resistance (AR) commonly found in bacteria able to survive in extreme acid conditions ^20–24^. In *E. coli*, the GAD system is a three components system: two glutamate decarboxylases, GadA and GadB, which use cytoplasmic free protons by converting glutamate into γ-aminobutyrate (GABA), and the glutamate/GABA antiporter GadC. When the pH is below 5.6, cytoplasmic GadB migrates near the inner membrane to maximize collaboration with transmembrane GadC ^25^. While *gadA* is independent in chromosomic location and *gadB* and *gadC* are organized in operon, the expression of both *gadA* and *gadBC* is transcriptionally regulated by RpoS, two AraC-like regulators GadX and GadW, and by effectors with two inhibitors, the cyclic AMP receptor protein and H-NS. H-NS and RpoS in particular determine the temporal expression, the former inhibiting *gadB* expression, whereas RpoS promotes the transcription of *gadB* once the stationary phase is reached ^26^.

To decipher and model fitness and behavior of *E. coli* O157:H7, synthetic microbial ecology approaches were used in structured food matrices where the complexity of the communities and the factors of influence are reduced to their minimum, but increased in their controllability ^27^. Such approaches have been used to describe how matrix parameters affect bacterial growth and morphodynamics of microcolonies ^16,28^. In a recent contribution, we have shown that the volume, distribution and sphericity of microcolonies of *E. coli* O157:H7 in hydrogel are dependent of the size of the inoculum, but also on the concentration of acids and NaCl, two environmental stresses frequently encountered in food products ^29^.

In this study, we took advantage of a hydrogel matrix to observe the local expression of *gadB* in *E. coli* O157:H7 cells in microcolonies using confocal laser scanning microscopy (CLSM). To explore the existence of patterns of expression in microcolonies, bacterial strains with a dual transcriptional fluorescent reporter system were engineered to monitor the spatial expression of *gadB* at the single cell scale. In order to relate the impact that phenotypic heterogeneity in microcolonies can have on community function, the survival of planktonically grown cells to a strongly acidic media mimicking the stomachal passage was further assessed and compared to cells grown or dispersed in hydrogels.

## 2) Material and methods

### Bacterial strains and culture conditions

From cryotubes stored at −80°C, the bacterial strain of *E. coli* (see genetic construction) was plated on Petri dishes with TSA (Tryptone Soya Agar, Oxoid, USA) and incubated overnight at 37°C. One bacterial colony was picked up and inoculated in TSB (Tryptone Soya Broth, Oxoid, England) before overnight incubation at 37°C under orbital shaking (200 rpm). When required, growth media were supplemented with chloramphenicol (Cm 25 μg/mL; EUROMEDEX, China). The strain *Lactococcus lactis* ssp. cremoris (Aerial N°2124) was incubated in the same conditions, but without antibiotic supplementation.

### Genetic construction

The *E. coli* O157:H7 CM454 ^30,31^ is the wild type strain in this study. We used a T7 polymerase (T7pol) amplification technique inspired from previous reports ^32,33^, where the cassette *T7pol::Cm*^*R*^ is inserted after the genetic sequence of the gene *gadB* using the Datsenko-Wanner ^34^ recombination technique (supplementary material Figure S1). Regions of identity were added at the ends of the cassette by the forward primer: 5’CCGAAACTGCAGGGTATTGCCCAACAGAACAGCTTTAAACATACCTGATAACAGGAGGTAAATAATGCACACGATTAACATCGC3’ and reverse primer: 5’AAATTGTCCCGAAACGGGTTCGTTTCGGACACCGTTACCGTTAAACATGGAGTTCTGAGGTCATTACTG3’. The correct insertion of *T7pol::Cm*^*R*^ in the construct was verified by PCR using the forward primer 5’GGAAGACTACAAAGCCTCCC3’ and reverse primer 5’ TATTCCTGTCGGAACCGCAC3’, for sequencing (Eurofins Genomics, Germany). Based on the sequence of the pHL40 plasmid ^32^, a new plasmid was synthetized (GeneArt, ThermoFisher Scientific, Germany) bearing the P_*T7pol*_*::GFPmut3::*T_*p7pol*_ as a GFP reporter but modified by insertion of P_*BBa_J23119*_*::mCherry2::*T_*BBa_B0062*_ (iGEM parts) for constitutive expression of a red fluorescent protein (RFP). This new plasmid, called pHL60, was transformed into competent *gadB::T7pol::Cm*^*R*^ bacterial cells. This system is an indirect reporter of *gadB* transcription as the transcriptional fusion of *gadB::T7 polymerase* allows an amplified production of GFP (GFPmut3) from pHL60 and normalization of the level of expression respective to the constitutive expression of the RFP (mCherry2) from the same plasmid, to minimize variations of the fluorescence associated with variations in the number of plasmids from one cell to another. To validate the genetic construction, the reporting planktonic expression of *gadB* was tested on six pH values from 4.5 to 7.0 using a microplate reader (Synergy H1, Biotek) (Supplementary material Figure S2).

### Transparent hydrogel matrices for fluorescent imaging

As previously described ^29^, the hydrogel matrices were obtained by mixing TSB with 0.50 % low melting point agarose (LMPA) (UltraPure Agarose, Invitrogen, USA). After boiling, the liquid LMPA at neutral pH (pH=7) was cooled down to 40°C to prevent thermal stress before the bacterial inoculum was added to obtain 10^4^ CFU/ml. When necessary, the medium was adjusted to acidic pH= 5 with HCl. After homogenizing and gentle stirring to avoid bubble formation, the inoculated gel matrix was immediately distributed in each well of a 96-well microtiter plate of microscopic grade (µClear, Greiner Bio-One, France). The microtiter plates were then incubated at 20°C and observed under CLSM after 96 hours of incubation.

All microscopic observations were performed with a Leica HCS-SP8 confocal laser scanning microscope (CLSM) at the INRAE MIMA2 imaging platform (https://doi.org/10.15454/1.5572348210007727E12). The GFP (GFPmut3; λ_ex_500; λ_em_513) and RFP (mCherry2; λ_ex_589; λ_em_610) were excited respectively with laser bands 488 nm and 561 nm. For Live/dead exploration, SYTO9 (λ_ex_485; λ_em_501) and IP (λ_ex_535; λ_em_617) were excited respectively with laser bands 488 nm and 561 nm. Observations were carried out with a water immersion 63x objective lens (numerical aperture of 1.20) for 184µm x 184µm fields. Bidirectional acquisition speed of 600 Hz allows a frame rate of 2.3 images per second. For 3D stack analysis, a 1 µm step between z levels was used. For each condition, a minimum of 60 stacks were acquired in over a dozen independent wells. Microscopic images were treated on IMARIS v9.64 (Bitplane, Switzerland) to generate sections and projections. Kymograms reporting the spatial analysis of *gadB* expression in microcolonies were performed using BiofilmQ v0.2.2 ^35^. BiofilmQ image segmentation was performed with a threshold value set at 0.1 with cubes of 1.8 µm (vox of 10). The absence of radial fluorescence gradients in microcolonies of *E. coli* O157:H7 constitutively expressing GFP was verified prior experiments with *gadB* expression (supplementary material Figure S3).

### Acidic digestion challenge

To test the ability of *E. coli* O157:H7 population to survive the strong acidic stress during the stomachal passage, 3 ml of planktonic cells (TSB), planktonic cells grown in TSB and then encased in gel matrix (TSB-LMPA) or gel-colonies cultures (LMPA) (72h, 20°C, pH=7 or pH=5) were transferred in 27 ml of NaCl 9g/L saline solution (control groups) or 27 ml of a saline solution adjusted with HCl (5 M) to a pH of 2. The cups were then incubated at 37°C for 4 hours under a 90-rpm shaking to simulate matrix digestion. All media were then homogenized to disperse bacteria (IKA Ultra-Turrax T25; Janke Kunkel) and the resulting suspensions were immediately plated on agar for enumeration and determination of the log reduction in CFU/ml before and after acidic treatment.

### Statistics

Graphics and ANOVA variance analysis were performed with Prism 9 (GraphPad; CA, USA). Differences were considered significant when *P*<0.05 with *P* being the critical probability associated with the Fisher test.

## 3) Results

### Spatial patterns of gadB expression in gel-microcolonies

Mean radius of microcolonies grown in neutral or acidic hydrogel showed little differences between neutral (28 µm) and acidic conditions (27 µm), the repartition of populations around these values was noted to not be statistically significant (Figure 1, obtained from measurement on 40 colonies, *P*>0.05). However, at pH 5, microcolonies appear more circular than at pH = 7 and they harbor at their edge a streamer population shedding from the colony core, forming a crown around it (Figure 2).

**Figure 1:**
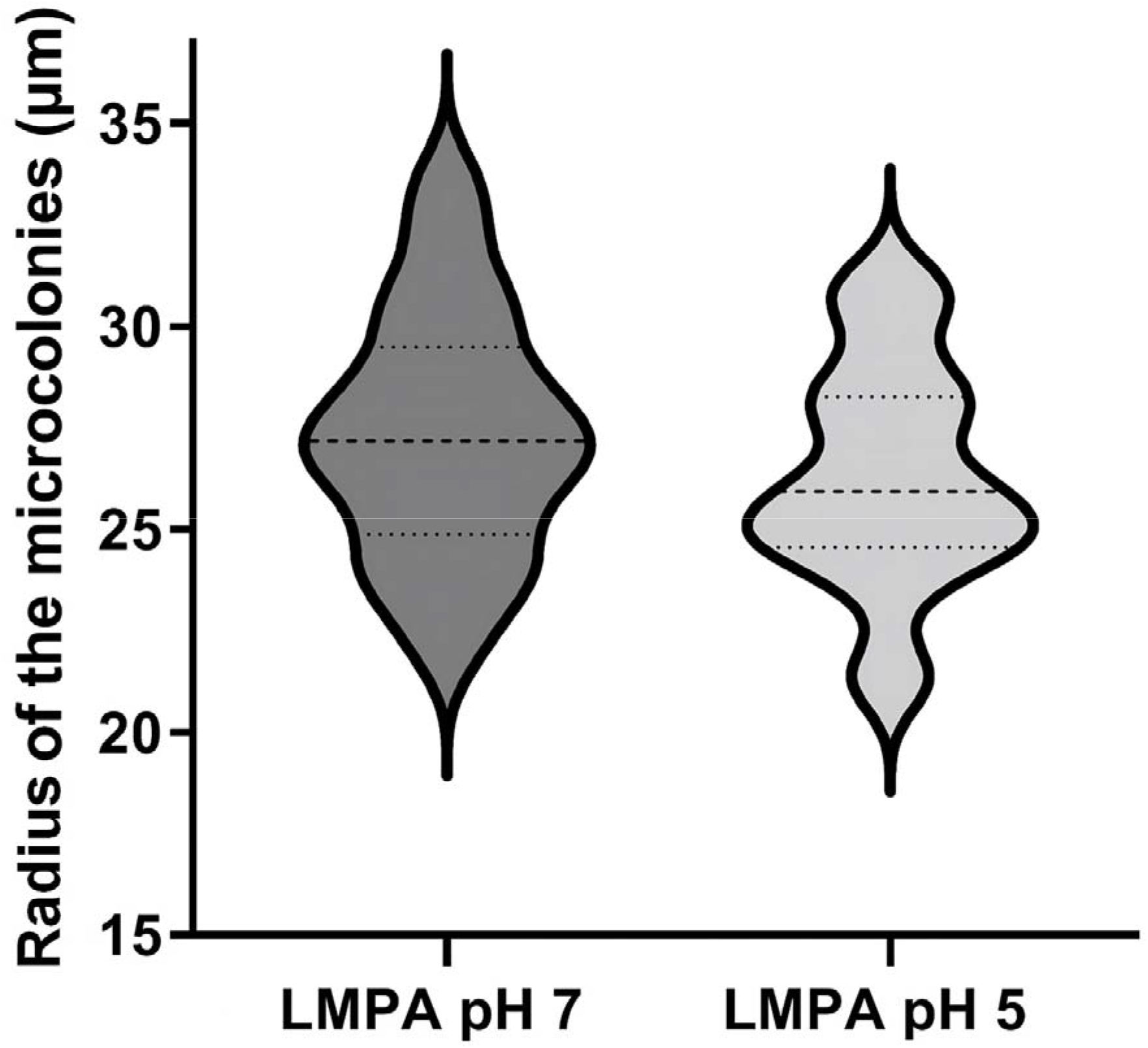
Radius of the microcolonies. Representation of the radius of the microcolonies in µm, the width of each figure represents the concentration of the number of values. For each case, the slashed line is the mean value of radius, and dotted lines delimit the 75 % probability interval. Radius values were calculated from 40 independent microcolonies.

**Figure 2:**
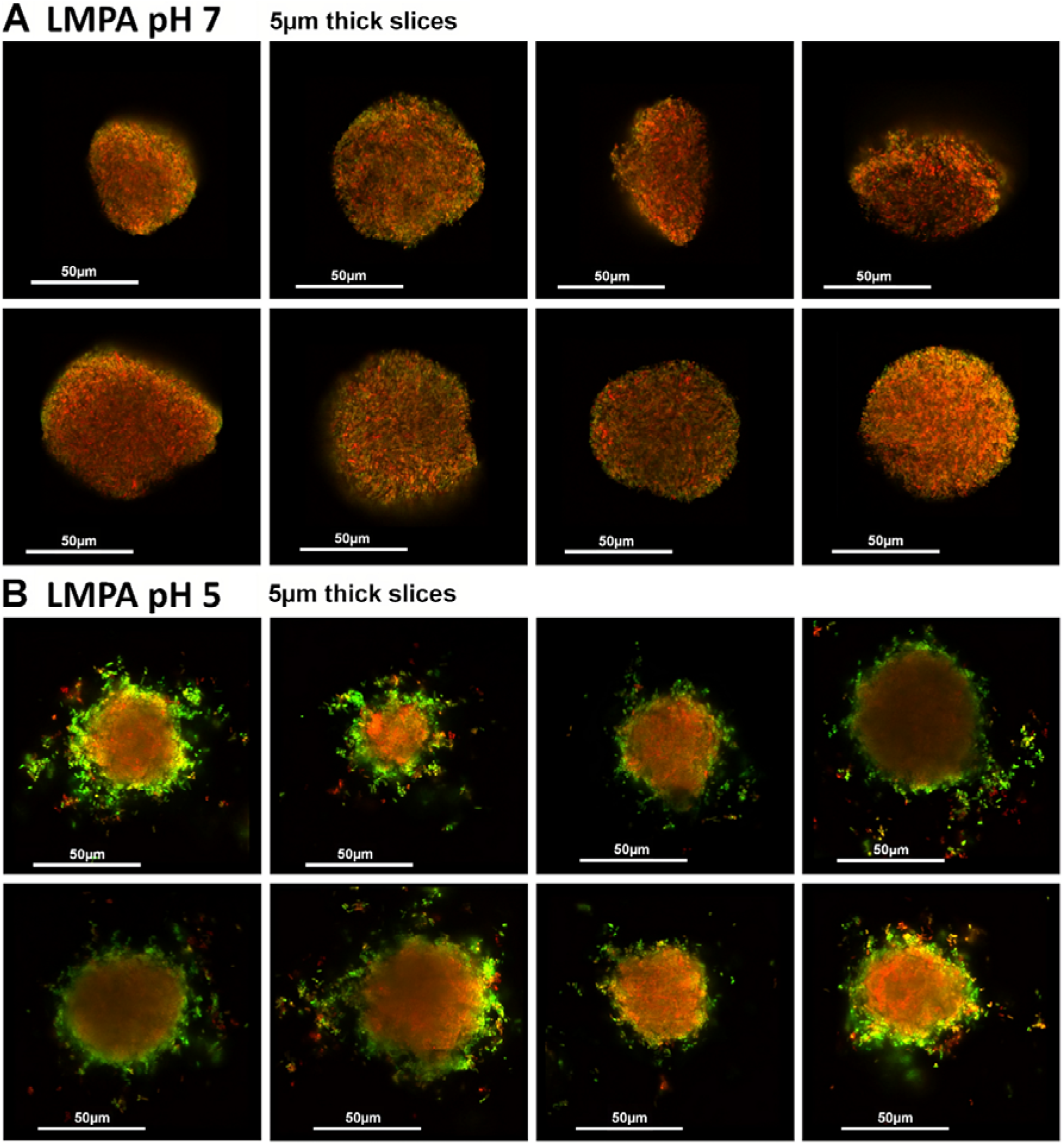
Representative images of *gadB* spatial patterns of expression for *E. coli* O157:H7 cultivated in neutral (pH=7) or acidic (pH=5) hydrogels. A series of 5µm slices of microcolonies is presented in control LMPA pH=7 (A) and the acid matrix LMPA pH=5 (B). The red fluorescence is constitutive and the green fluorescence is expressed as a function of *gadB* transcription. Supplementary material Figure S3 presents the same representation for a control constitutive GFP expression in both conditions.

Microscopic observations of the fluorescence reporting the expression of *gadB* in cells inside microcolonies (Figure 2) show radically different patterns between the two hydrogels. In neutral pH conditions, the gene is expressed at a basal level throughout the whole microcolony with no specific spatial arrangement (Figure 2A, supplementary material Figure S4). By contrast, the expression of *gadB* is strongly overexpressed in the periphery of the colonies formed in acidic hydrogel (Figure 2B, supplementary material Figure S4). Those qualitative observations were reinforced by a quantitative exploration of the radial distribution of *gadB* expression (Figure 3). For both hydrogels, the genetic expression is monitored by the green fluorescent intensity normalized with the red constitutive fluorescent intensity. 3D kymographs integrating 40 independent microcolonies (X-axis) for each condition represent in color code *gadB* transcription from the center of the microcolony (Y-axis, d_CM_=0 µm) to the edges of the colonies and beyond. Where in neutral pH hydrogel *gadB* expression is low and almost constant over the radius of the microcolonies (Figure 3A), acidic hydrogels present a sharp band of *gadB* over-expression in between 25 and 30 µm from the center of the microcolonies (Figure 3B). This is consistent with the observed spatial expression as the radius of the microcolonies is 27-28 µm (±5 µm) in these experimental conditions. Interestingly, when microcolonies merge as they grow, they behave like a single colony in regard to the peripheral *gadB* spatial expression. Similarly, if two microcolonies are in near contact, the two sides facing each other do not present an over-expression of *gadB* or the shedding of single cells visible in other areas of the periphery (supplementary material Figure S5).

**Figure 3:**
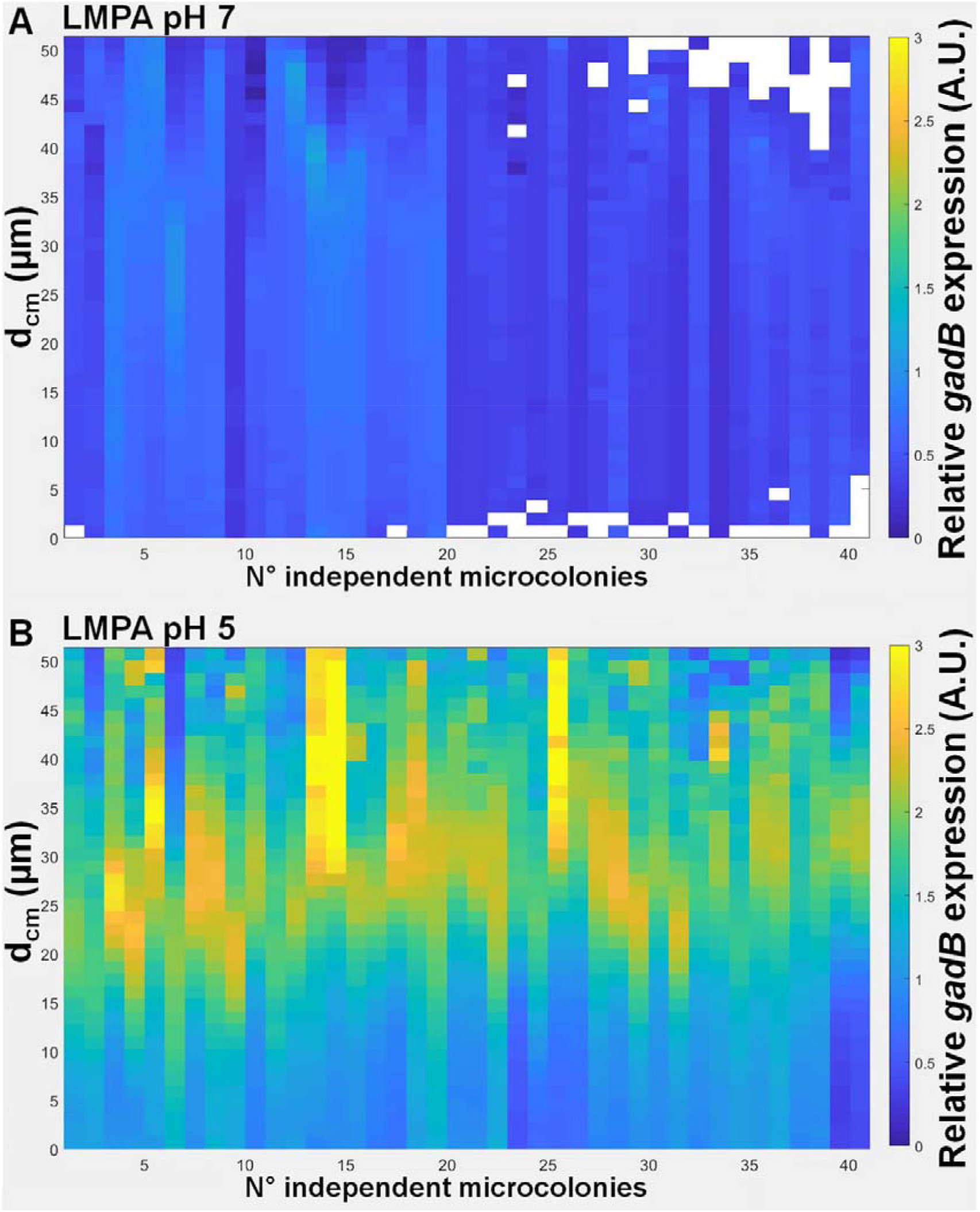
Relative expression of *gadB* in function of the distance from the center of microcolony. In this representation, the relative *gadB* expression is expressed as the ratio of green fluorescence intensity over red fluorescent intensity (GFP reporter/RFP constitutive) in function of the distance from the center of the microcolony (d_CM_) for forty microcolonies grown in neutral (A) or acidic hydrogels (B).

Control experiments performed with a constitutive expression of the green fluorescence protein did not show the spatialization of the expression as presented above (supplementary material Figure S3). A time-course microscopic analysis allows the observation of *gadB* expression spatialization in acidic hydrogels as early as microcolonies become visible under the microscope, ∼48h after inoculation (data not shown). Finally, the spatial repartition of dead cells in gel-microcolonies as shown by live/dead fluorescent staining indicated a random distribution of the red dead cells, with no preferential localization in the microcolonies associated with cells expressing *gadB* (supplementary material Figure S6).

Following those results, *E. coli* O157:H7 was then cultured in the presence of *Lactococcus lactis* ssp. *cremoris* (Figure 4). The ability of *L. lactis* to produce *in-situ* lactic acid is used to replicate the natural acidification of food matrix containing *L. lactis* (cheese), or where a progressive accumulation of lactic acid occurs (meat). Exploration of the hydrogel was performed 72-96h after inoculation. Contrary to observations where lactic acid is added in the hydrogel preparation ^29^, microcolonies of *E. coli* O157:H7 are present and possess the same morphology as seen in mono-cultures in the acidic condition (Figure 4A). Close up on 50 µm thick slices of microcolonies of *E. coli* O157:H7 in close proximity to microcolonies of *L. Lactis* clearly shows the same spatial patterns of *gadB* expression as previously encountered for the mono-cultures in acidic media (Figure 4B).

**Figure 4:**
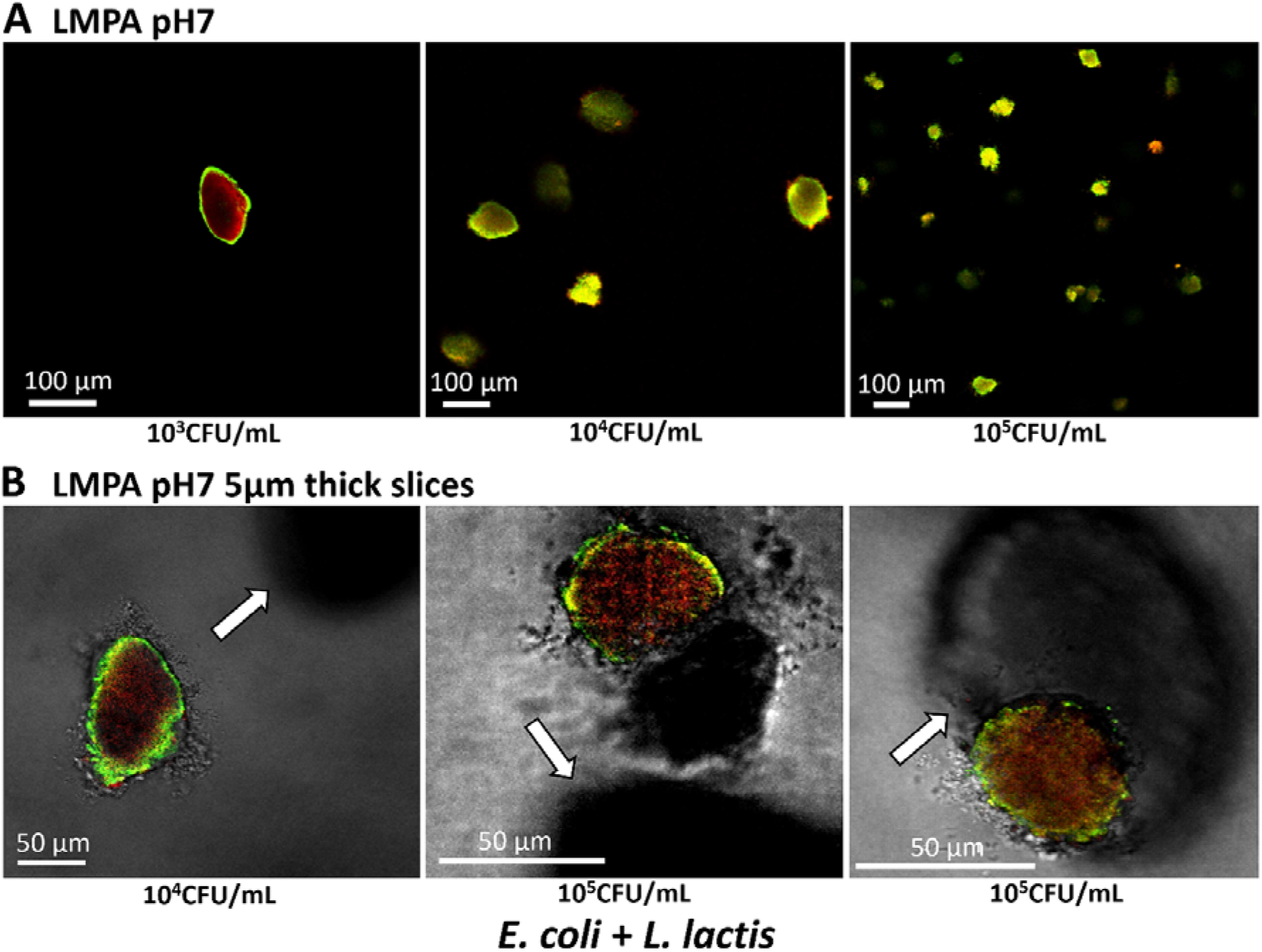
Spatial patterns of expression of *gadB* for *E. coli* O157:H7 in the presence of *Lactococcus lactis* ssp. *cremoris*, cultivated at neutral pH in hydrogels. (A) Visualization at 96h of microcolonies of *E. coli* O157:H7 co-inoculated from 10^3^CFU/ml to 10^5^CFU/ml, with *L. lactis* inoculated at 10^3^CFU/ml, in a hydrogel at neutral pH. The red fluorescence is constitutive and the green fluorescence is expressed as a function of *gadB* transcription. (B) Microcolonies of *E. coli* O157:H7 close or in contact with *L. lactis* microcolonies. The two rightmost pictures were taken at 96h and the one on the left at 72h. *L. lactis* microcolonies are visualized in the bottom images thanks to the transmission detection (indicated by white arrows).

### Increased survival to acidic stomachal stress of E. coli grown in gel-microcolonies

As the capacity of survival to stomachal acidic stress of *E. coli* O157:H7 populations is of interest for public health safety, the acid resistance of bacteria grown planktonically or in hydrogel matrices was evaluated by enumeration on agar after acid stress.

Cultures of *E. coli* O157:H7 adjusted to 10^4^ CFU/ml were incubated at neutral (pH=7) or acid pH (pH=5) in either TSB or LMPA. After 96h of incubation at 20°C, the populations reached values of log CFU/ml of 9.4/9.6 in TSB at pH neutral/acid and 9.5/8.6 in LMPA at pH neutral/acid.

Bacteria grown in planktonic conditions (TSB) were highly sensitive to the 4-hours exposition to pH=2 with a total loss of the culturable population (9 log reduction) (Figure 5). In contrast, cells grown as spatially organized colonies in LMPA for 96h presented a statistically high tolerance to this strong acidic stress (*P*<0.05). The best tolerance was observed for the microcolonies incubated at a pH of 5 with a log reduction as low as 0.29 log CFU/ml, statistically significantly lower than the reduction observed for microcolonies incubated at neutral pH, where the log reduction is 0.78 CFU/ml (*P*<0.05). To test for interferences of the hydrogel to bacterial acidic stress, planktonic populations cultivated in TSB were encased in LMPA just before the survival test. Log reduction of these control planktonic populations suspended in LMPA presented significant similar sensitivity than planktonic TSB culture (*P*>0.05), indicating no buffering effect of agarose to stomachal acidic stress.

**Figure 5:**
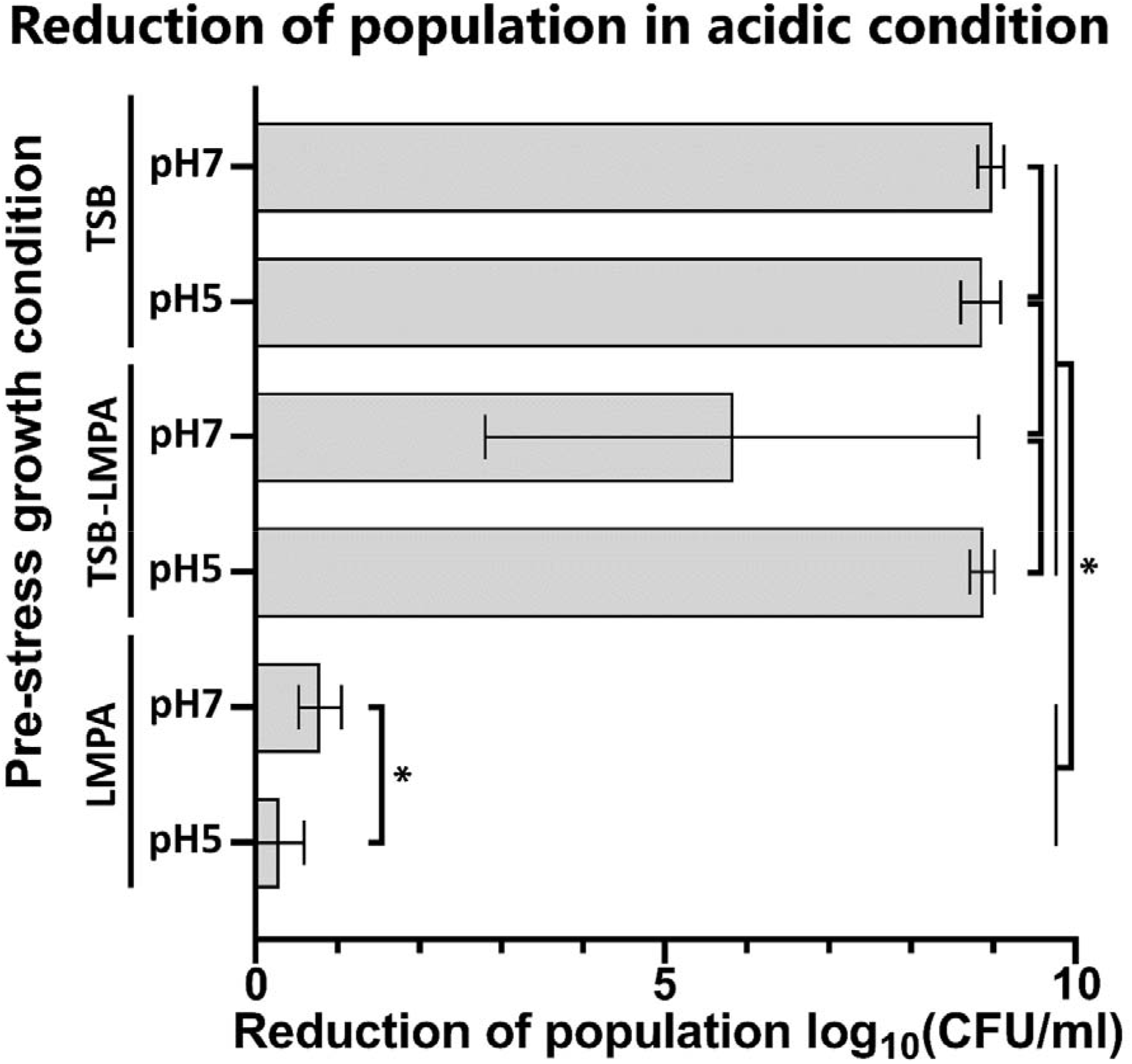
Population log-reduction of planktonic cultures and hydrogel microcolonies of *E. coli* O157:H7 upon acidic exposure. Bacterial cells were exposed for 4 hours at a pH of 2 (HCl). Representation of the mean reduction of population log between the control and survival groups. Bacteria incubated and tested in planktonic are on the top (TSB), those incubated in planktonic but encased in hydrogels before the test are on the middle (TSB-LMPA), and the results for populations incubated and tested in hydrogels are on the bottom (LMPA). A star indicates a significant difference between values (*P*<0.05). Data resulted from at least six biological replicates.

## 4) Discussion

The behavior of microbial population in laboratory liquid growth media can strongly deviate from what is observed in real solid food matrices ^18,36–40^. The environmental heterogeneity of structured media is listed as one of the four main causes of cellular variation, among genetic variation, aging and stochasticity of gene expression ^41^. As such, it can trigger a large diversity of phenotypic cell expression in the same biotope, promoting the cohabitation of cells with different spectra of behaviors, such as stress response or virulence ^42,43^. Structured food are not an optimal medium from an exploration perspective as the opacity of numerous food matrices prevents live imaging and microscopic approaches. To overcome these limitations, several studies take advantage of synthetic hydrogelled systems to simplify and control the parameters of growth of embedded bacteria. Here, low melting point agarose is used as a gelling agent in which cells can be dispersed without thermal stress and with tunable textures and media compositions. Thus, in a recent contribution, we have been able to mimic the texture of various food environments such as grounded meat or cheese ^29^. The experiments presented in this work show that the morphology of microcolonies in the media complemented with HCl are different from the neutral pH control, in particular bacteria are shedding from the periphery of the microcolonies. This effect can be explained by a combined action of relaxed gel structures due to low pH and the higher motility of *E. coli* O157:H7 when acidic conditions are encountered ^29,44^.

In this study, we observed a clear overexpression of *gadB* for a subpopulation of cells localized in the periphery of microcolonies formed in acidic hydrogels. This gene is overexpressed at levels 2-3 times higher than in neutral conditions, which is in accordance with results obtained in planktonic conditions (Supplementary material Figure S2). We confirmed that this spatialization was neither associated with dead cells (supplementary material Figure S6), nor a limitation of oxygen for GFP maturation in the center of the microcolony (supplementary material Figure S3). The use of a single plasmid bearing both the genes for the constitutive and induced fluorescence means that, at the image analysis step, we prevented bias due to differences in plasmid copy numbers or differences in coloration from a mix of dye/genetic reporters ^45^. The use of two lasers with different properties of matrix penetration could lead to a bias in the Z axis, but 3D analysis of all the cells in a microcolony reduces the bias as any offset at the bottom of the agglomerate is compensated by an opposite offset at the top.

Spatial patterns of genetic expression were previously reported for other genes in other bacterial species in surface biofilms either on solid or liquid, such as localized expression of *E. coli* sigma factors and type 1 pili, as well as *Pseudomonas aeruginosa* β-lactamase in biofilms ^46–48^. To our knowledge, such patterns of gene expression were never reported in food or hydrogel matrices. Last experiments of co-cultures of *E. coli* O157:H7 and *L. lactis* demonstrate that the pattern of *gadB* expression could naturally occur in food matrix through a progressive accumulation and diffusion of lactic acid in the media, such as in cheese or meat products.

Then, we explored the consequences of growing populations in a structured media in regards of survival to an exposition to low pH media. Results showed that, regardless of the initial pH, populations incubated in a semi-solid media have a better tolerance to acid stress than those grown in liquid broths, where no surviving cells were detected. This underlines limitations in modeling food-borne pathogens behavior in food from data obtained in liquid conditions, as previously shown in other studies^18,36,37,49^.

It has been suggested from other studies that the components of the matrix could have a buffer effect that protects embedded cells by preventing the drop in pH. Our tests show that planktonic populations dispersed in hydrogel did not present the same survival fitness that those cultivated as microcolonies in the same hydrogel. The hypothesis of a buffer effect due to hydrogel interference was tested for bacteria incubated in hydrogel and it was statistically rejected. This is supported by another study in a gelified dairy matrix, where food related bacteria were dispersed without incubation in the gel before application of the acidic stress. It was reported that no protective effect existed compared to the same conditions in liquid media ^50^.

A parameter that could explain this difference of survival between the populations incubated or not in the hydrogels would be the spatial organization of cells. The hydrogels showed evidence of deliquescence such as unraveling of filaments and loss of stiffness but maintained enough structural integrity to ensure the microcolonies did not disperse. The ability of spatially organized populations of bacterial cells to better survive acid stress was described for pathogenic bacteria but also for auxiliary microbiota and probiotics, such as *Lactobacillus* strains ^51,52^. The bacteria could secrete extracellular polymeric substances (EPS) when grown in communities as described in biofilms ^53,54^. For *E. coli* O157:H7, tolerance of the bacteria to low pH could further involve the DNA binding protein Dps which is known to enhance survivability when local nutrients are exhausted ^55,56^.

From this study in a hydrogel matrix, *gadB* appeared to be more expressed at the periphery of the *E. coli* O157:H7 microcolonies in acidic conditions. This correlated with an increased tolerance to the type of acid stress that can be encountered by bacterial cells after ingestion of food. These findings are of interest for public health as they underline possible differences between liquid and solid food products on the infectious dose and bacterial virulence. In this case, the tolerance to acidity could mean an increase in the bacterial load that can survive in the digestive system as well as a phenotype more likely to colonize the gut lining. In order to alleviate public health issues, differences in bacterial behavior in planktonic conditions versus microcolonies could be considered when integrating phenotypic heterogeneity in risk assessment. Modeling pathogens growth and survival should take in account the gelled environments where the spatialization of genetic expression and its resulting populational effects could deeply affect pathogens behavior and virulence after ingestion.

## Abbreviations

CLSM: stands for “ Confocal Laser Scanning Microscopy”
LMPA: stands for “ Low Melting Point Agarose”

## Declaration of competitive interest

None

## Acknowledgments

This work was supported in part by INRAE and ANR (Agence National de la Recherche) PathoFood project (n°ANR-17-CE21-0002). The partners of the ANR PathoFood are acknowledged for fruitful discussions about *L. monocytogenes* and *E. coli* O157:H7 behavior in food matrices. Cédric Saint Martin was a PhD Research Fellow granted by the ANR PathoFood. Julien Deschamps (INRAE) is warmly acknowledged for assistance with Confocal Laser Scanning Microscopy at the Mima2 imaging platform (https://doi.org/10.15454/1.5572348210007727E12).

## Author contributions

CSM, RB and MD, conceptualized the overarching aims of the research study. CSM, NC, MD, MG, AC, GJ, FDB, SL, RB and MD conceived and designed the experiments. CSM, NC, MD, MG and AC performed the experiments and data acquisition. CSM, NC, MD, AC, GJ, FDB, SL, RB and MD analyzed and interpreted the data. RB and MD had management as well as coordination responsibility for the execution of the research work. RB and MD contributed to the acquisition of the financial supports and resources leading to this publication. CSM, NC, MD, MG, AC, GJ, FDB, SL, RB and MD wrote the article, including drafting and revising critically the manuscript for important intellectual content. All authors have declared no competing interests.

## Supplementary materials

**Figure S1:**
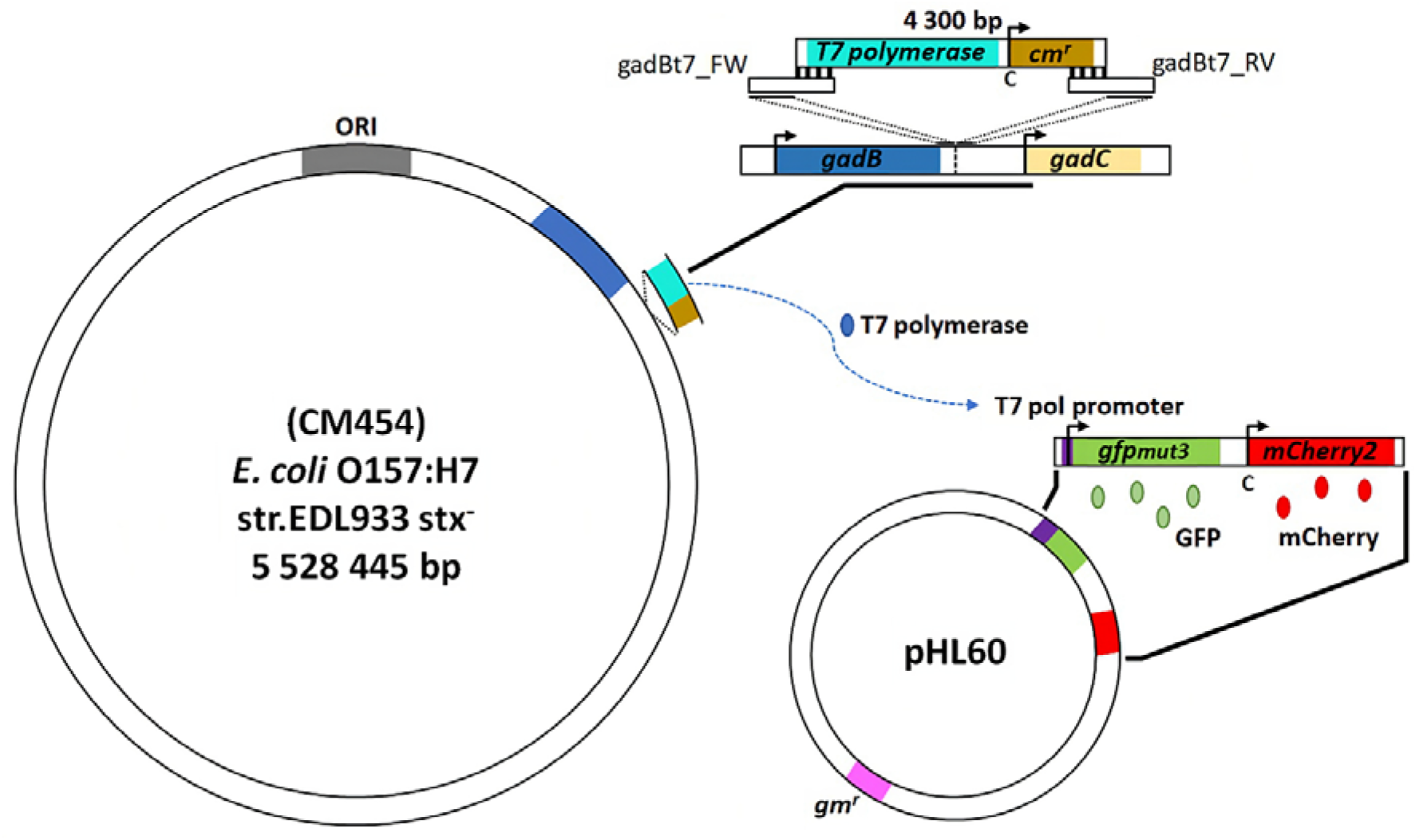
Strategy of the genetic constructions. (A) chromosome of *E. coli* O157:H7 with a zoom on the region of the insert and how the different CDS are present in this situation. The cassette bearing the *T7 polymerase* (in turquoise) and resistance gene *cm*^*r*^ (in brown) is elongated by PCR with two primers (gadBt7_FW and gadBt7_RV) so that a homology sequence exists with insertion site. This site is located between the gene interest *gadB* (in blue) and *gadC* (in yellow). On the right the low copy plasmid pHL60 bearing the fluorophore encoding genes is represented. It contains the constitutively expressed *mCherry2* (in red) sequence and *gfpmut3* (in green) under the control of the *T7 pol promoter* (in purple), as well as a gentamicin resistance gene *gm*^*r*^ (in pink).

**Figure S2:**
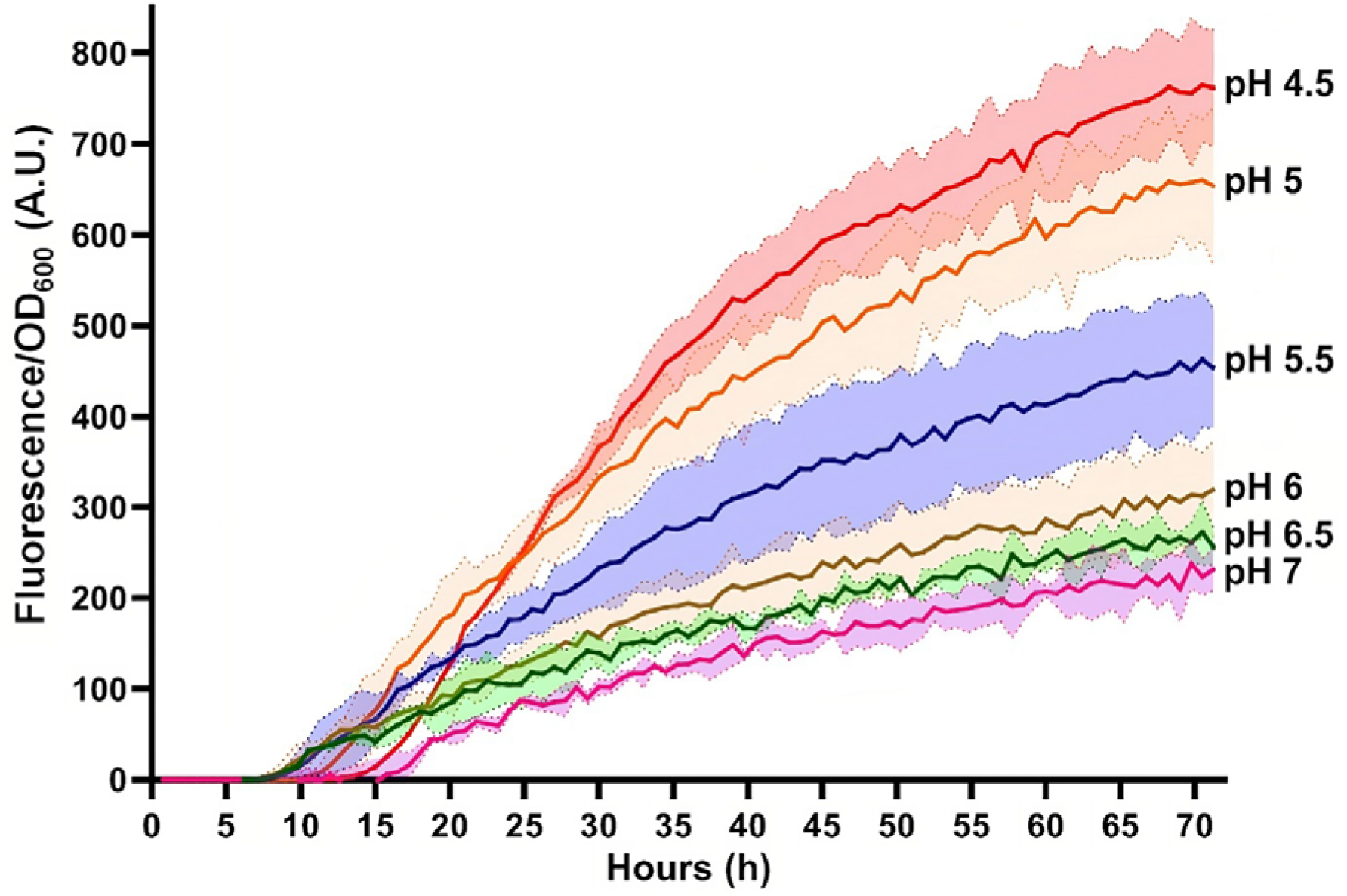
Relative expression of *gadB* in a population in function of time and under different acidity levels. The fluorescence of planktonic populations of *E. coli* O157:H7 *gadB::GFP* are represented in function of the OD_600_. For each curve representing the mean signal over time, the standard deviation of values is shown as an area of lighter color. These curves show that detected fluorescence becomes incrementally brighter with decreasing pHs. The rise in expressed fluorescence is not linear with greater leaps of intensity below pH=6.0. Points over time show that fluorescence intensity at pH 4.5 can be 3-4 times higher than reported values for the pH=7.0 control. The lag time before the green fluorescence is detected is high at pH 7.0 (15 hours) compared to pH=5.5 where it starts after eight hours. In media pH=5.0 and pH=4.5, the lag time becomes progressively longer (12 then 14 hours).

**Figure S3:**
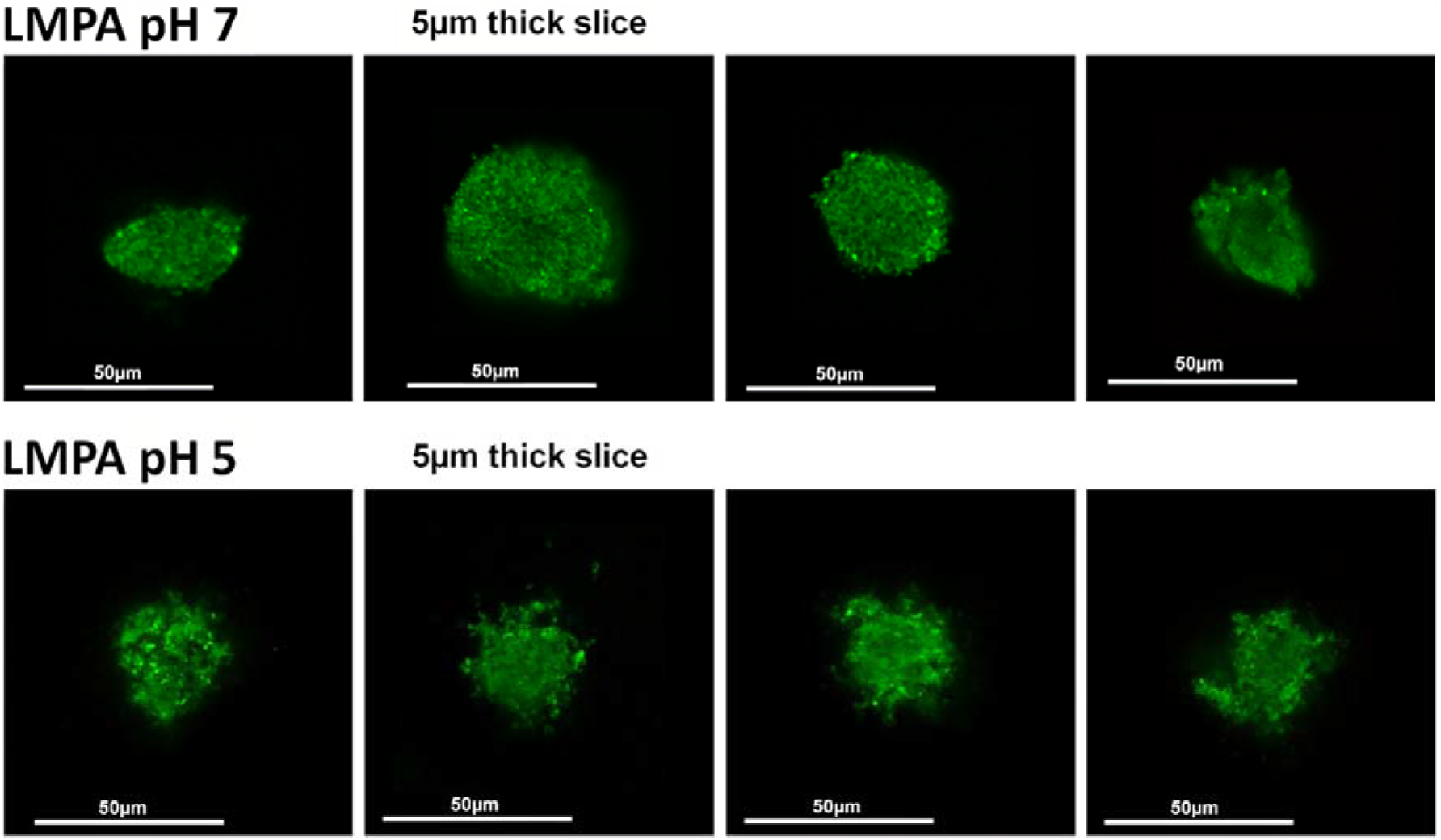
Qualitative presentation of the constitutive GFP spatial expression in microcolonies at pH=7 or pH=5. A control group with a constitutive GFP to verify that at pH=7 (up) and/or pH=5 (down) the fluorescence of the green fluorescence is not already spatialized. Each image is a 5µm slice of a stack.

**Figure S4:**
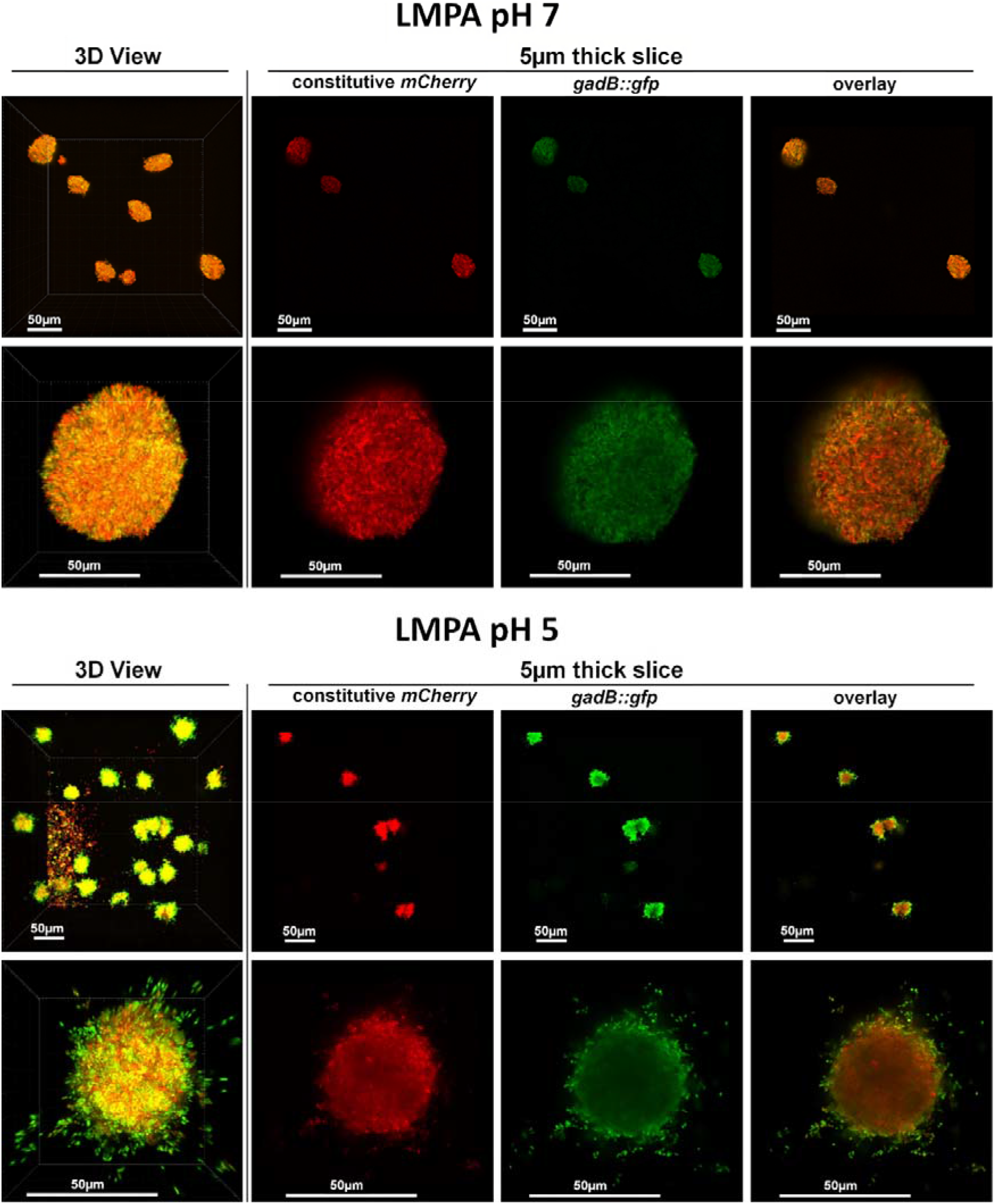
Qualitative presentation of *gadB* spatial expression in microcolonies at pH=7 or pH=5. Microscopic observations of the expression of *gadB* in the control LMPA pH=7 (top) and in the acid matrix LMPA pH=5 (bottom). For both groups, the 2 successive rows use a 40X then a 63X objective to present a collection of microcolonies and a close observation of a single example. Each image is shown as a 3-dimensional representation (leftmost column) and in each case a series of 5 µm slices show both fluorescent channels (middles columns) and the overlay (Rightmost column). The first row of each condition was taken with a 40x air objective (numerical aperture = 0.85) to explore a 290 µm x 290 µm fields.

**Figure S5:**
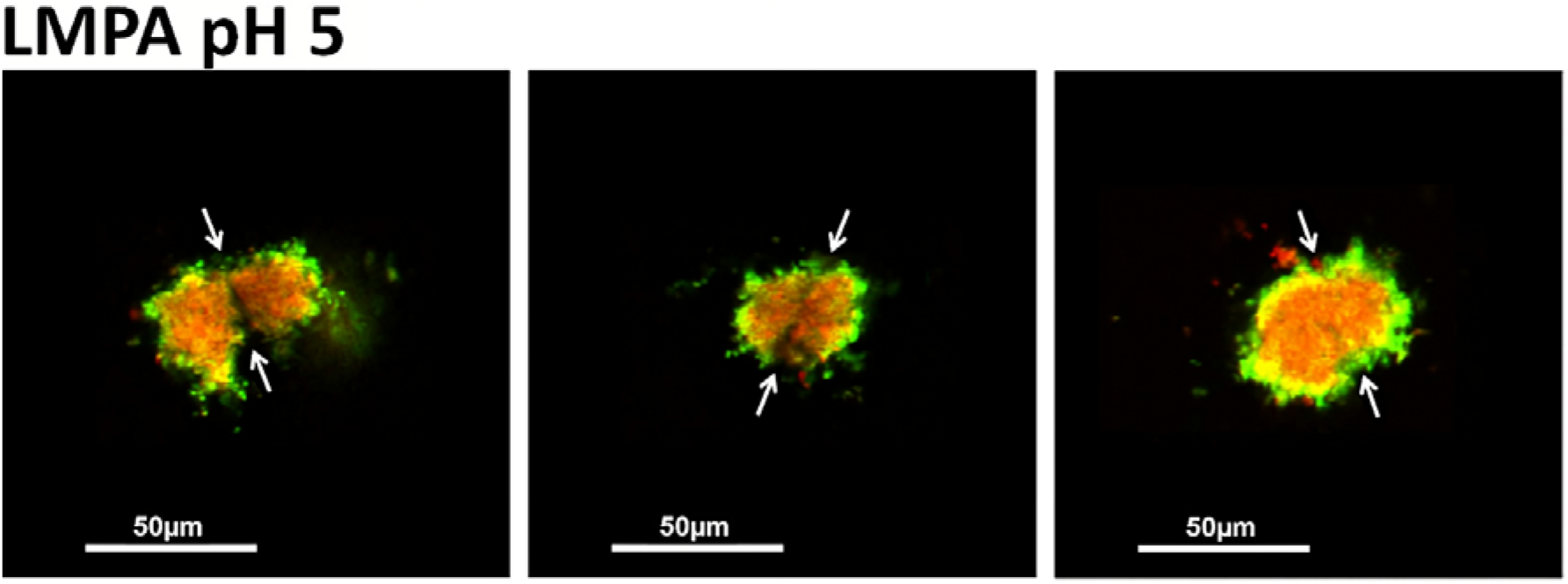
The case of joint microcolonies. Examples where two microcolonies of *E. coli* O157:H7 are touching or merging in LMPA pH=5. White arrows indicate the separation between each microcolonies.

**Figure S6:**
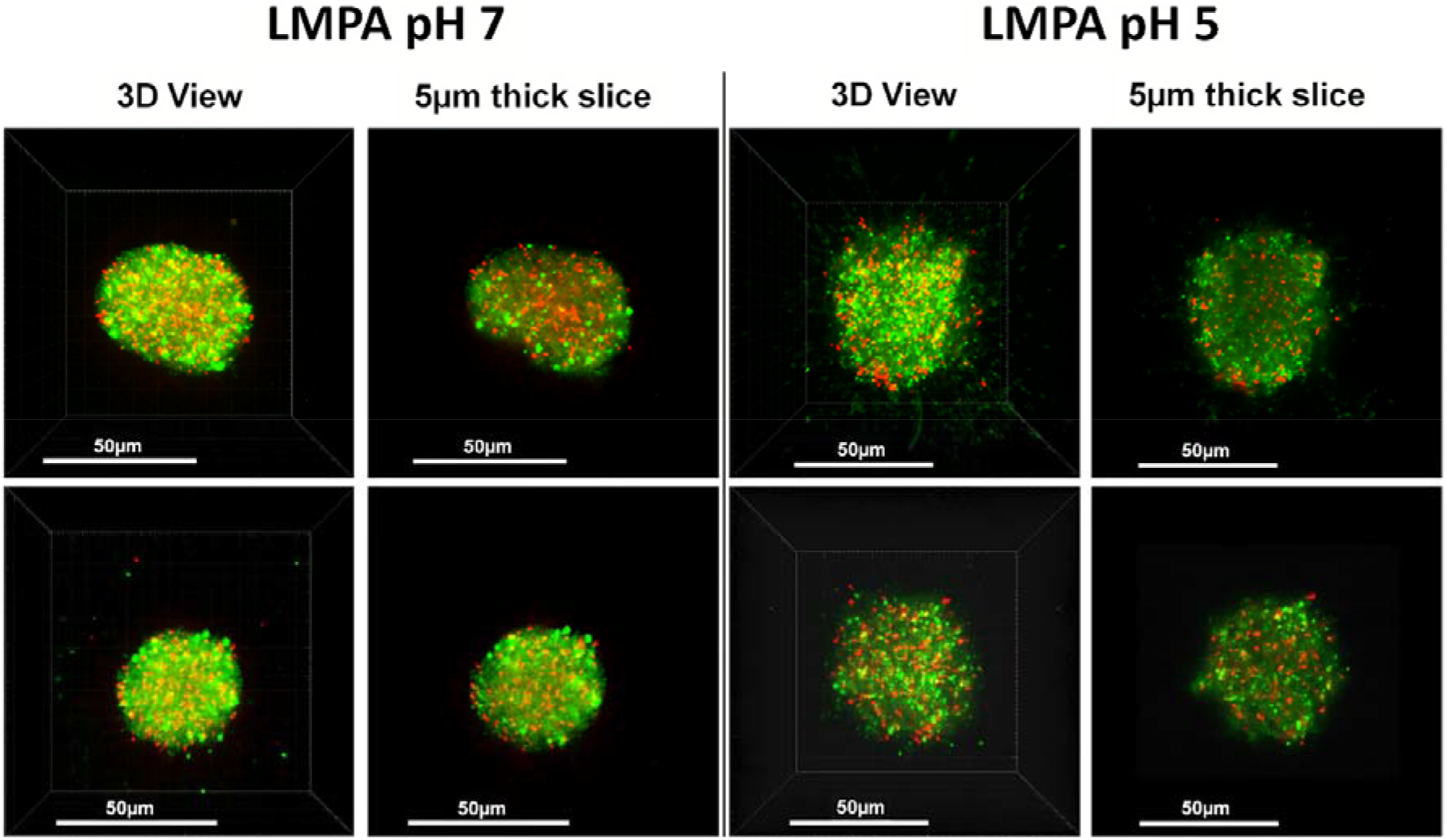
Live/Dead staining in the gel microcolonies. Bacterial cells in microcolonies grown in neutral (left side) or acidic (right side) were labeled with the cell impermeant propidium iodine (red, dead cells) and the cell permeant SYTO 9 (Green, all cells).

